# Ilaprazole and other novel prazole-based compounds that bind Tsg101 inhibit viral budding of HSV-1/2 and HIV from cells

**DOI:** 10.1101/2020.05.04.075036

**Authors:** Jonathan Leis, Chi-Hao Luan, James E. Audia, Sara F. Dunne, Carissa M. Heath

## Abstract

In many enveloped virus families, including HIV and HSV, a crucial, yet unexploited, step in the viral life cycle is releasing particles from the infected cell membranes. This release process is mediated by host ESCRT complex proteins, which is recruited by viral structural proteins and provides the mechanical means for membrane scission and subsequent viral budding. The prazole drug, tenatoprazole, was previously shown to bind to ESCRT complex member Tsg101 and quantitatively block the release of infectious HIV-1 from cells in culture. In this report we show that tenatoprazole and a related prazole drug, ilaprazole, effectively block infectious Herpes Simplex Virus (HSV)-1/2 release from Vero cells in culture. By electron microscopy, we found that both prazole drugs block the release of HSV particles from the cell nuclear membrane resulting in their accumulation in the nucleus. Ilaprazole also quantitatively blocks the release of HIV-1 from 293T cells with an EC_50_ of 0.8 μM, which is more potent than tenatoprazole. Finally, we synthesized and tested multiple novel prazole-based analogs that demonstrate both binding to Tsg101 and inhibition of viral egress in the nanomolar range of HIV-1 from 293T cells. Our results indicate that prazole-based compounds may represent a class of drugs with potential to be broad-spectrum antiviral agents against multiple enveloped viruses, by interrupting cellular Tsg101 interaction with maturing virus, thus blocking the budding process that releases particles from the cell.

**Importance:** These results provide the basis for the development of drugs that target enveloped virus budding that can be used ultimately to control multiple virus infections in humans.

## Introduction

The advent of antibiotics had a major impact on controlling bacterial infections in patients worldwide, with a single drug being used to treat multiple infections. Unfortunately, antivirals have not had the same success. There are many contributing factors to this shortcoming, foremost the fact that few mechanisms are shared by different viruses limits targets for a broad-spectrum antiviral. Consequently, approved antivirals generally act against individual rather than groups of viruses, limiting a single drug’s potential. However, this may change with the finding that multiple classes of enveloped viruses share the same budding mechanism that relies on host-cell endosomal sorting complex required for transport (ESCRT) proteins (1, 2). An inhibitor of this pathway could represent a potential broad-spectrum antiviral and have a positive impact on our ability to treat multiple enveloped virus infections with a single therapeutic agent.

Enveloped viruses bud from the host cell membranes and use the acquired lipid layer as a protective coat that also contains the glycoproteins required for infection of other cells. Enveloped viruses do not encode the machinery needed for budding and must recruit host-cell proteins to bud from cells. In HIV, ESCRT proteins are recruited to virus budding complexes through an interaction between the L-domain (PTAPP motifs) in virus structural proteins (3–7) with cellular protein Tsg101 (Tumor susceptibility gene 101), a homolog of the E2 ubiquitin conjugating enzyme and a member of the ESCRT-I complex (6, 8–11). Tsg101 recruits the cellular ESCRT-III complex, which provides the mechanical means for viruses to passage through cell membranes to be released (10, 12–19). Another enveloped virus family member, herpes simplex virus, HSV, assembles particles in the nucleus and relies on ESCRT proteins for passage through the nuclear membrane (20–23). Thus, this pathway may present a common target for treating multiple virus infections.

In support of targeting this pathway, a recent seminal discovery in our lab established that an interferon-induced protein, Interferon Stimulated Gene 15 (ISG15), specifically targets the ESCRT proteins in budding complexes to block the release of viruses (1, 24–26). This indicates that the human immune system evolved to target the ESCRT pathway to control infections and supports that this is a natural target. Another group identified single-nucleotide polymorphic sites in the 5’ region of Tsg101, located at positions −183 and +181 relative to the translation start signal, which affect the rate of AIDS progression among Caucasians (27). These data support the hypothesis that variation in Tsg101 affects efficiency of Tsg101-mediated release of viral particles from infected cells, altering plasma viral load levels and subsequent disease progression. Taken together, these investigations indicate that Tsg101 and ESCRT proteins present a natural antiviral target.

Currently the prazole family of drugs is best known for their role as proton pump inhibitors (PPIs) and a few, namely omeprazole (Prilosec), esomeprazole (Nexium) and ilaprazole (Adiza, Noltec, Yi Li An), are marketed to control symptoms of gastroesophageal reflux disease (GERD) in either the US or abroad. PPIs form a covalent bond with the active site of proton pumps, inhibiting their ability to acidify the stomach and reducing symptoms associated with over-acidification (28). Recent reports indicate that drugs from the prazole family, including tenatoprazole and esomeprazole, form a disulfide linkage to Tsg101, which results in blocking the release of HIV-1 from cells in culture (5). Other groups recently reported that prazoles may have potential as an antiviral therapeutic in HSV and SARS-CoV2 in combination with acyclovir or remdesivir, respectively (29, 30). However, the prazole compound used in these studies, omeprazole, was not potent enough to be predicted to have a therapeutic effect in vivo. This highlights a gap in the ability for current prazoles to have therapeutic potential, and the need for further research on prazoles as antivirals.

In the present manuscript, we demonstrate that multiple prazole drugs block the budding of HSV-1 and HSV-2 from Vero cells in culture, strengthening the case for the broad-spectrum potential of this mechanism/drugs. Most notably, we identified one prazole drug, ilaprazole, which blocks the release of both HIV-1 and HSV-1/2 from cells at an efficiency more potent than reported for tenatoprazole. Ilaprazole acts in the low μM range without detectable cell toxicity at inhibitory concentrations. Additionally, we designed and synthesized novel prazole analogs that act in the nanomolar range to block virus release, a major step forward in creating a VBI that can be brought to the clinic. To further define the mechanism of action of these prazole drugs on HSV infections, we identified the site of blockage of herpes virus release, which appears to be different from HIV-1. While the blockage to HIV-1 particle release is at the outer cell membrane (5), the prazole drugs appear to first block the passage of the herpes virus through the nuclear membrane. This prevents particles being released into the cytosol, where maturation of their envelope membrane occurs to produce infectious virus. With the prazole-based inhibitors being effective in both HIV and HSV, targeting Tsg101 could lead to a broad-spectrum antiviral therapy.

## Results

### Identification of prazole compounds that bind the UEV ubiquitin-binding domain of Tsg101

We screened chemical compounds using a fluorescence thermal shift (FTS) assay (31, 32) to identify small molecules that bind directly to a truncated form of Tsg101 (amino acids 1-145) which contains the Ubiquitin E2 variant (UEV) ubiquitin-binding domain (Fig. 1). This truncated Tsg101, called Tsg101-UEV, was used because full-length Tsg101 has significant solubility issues in aqueous solution. Tsg101 is an adaptor protein and thus lacks a readily deployable functional assay, making FTS a tractable approach to identify interacting compounds. FTS monitors protein thermal denaturation using SYPRO^®^ Orange, a dye which fluoresces when bound to hydrophobic surfaces, which allows monitoring of the changes in hydrophobic surface exposure during protein denaturation (31). Since ligand binding affects protein thermal stability, it can be detected through modulation of protein thermal denaturation (melting) as a shift in melting temperature (T_m_). Tsg101-UEV has a well-defined melting curve suitable for FTS. We used the FTS assay to identify compounds that bind to Tsg101-UEV.

**Figure 1.**
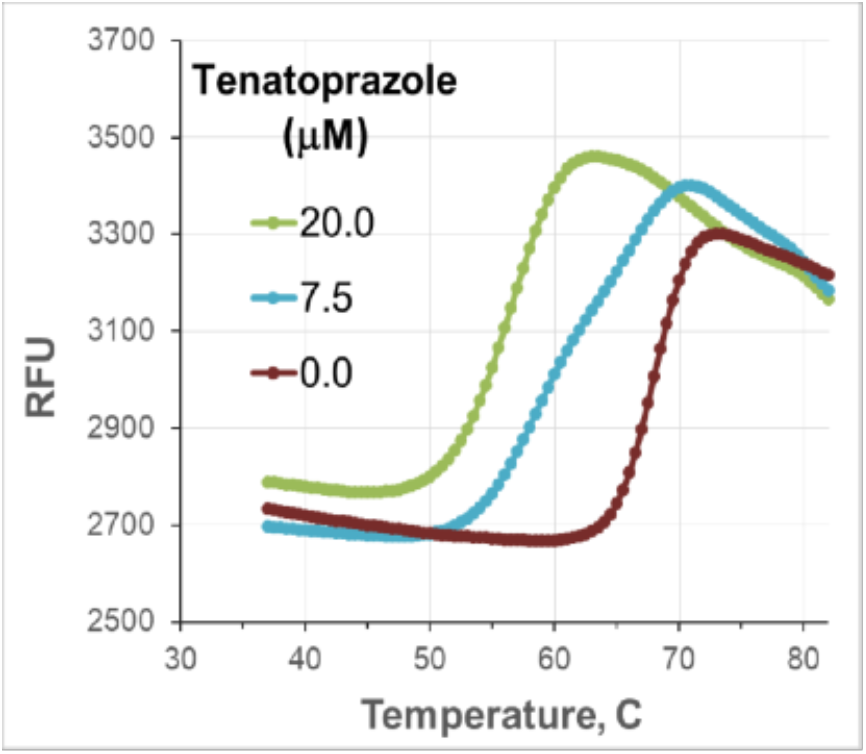
Thermal shift data of Tsg101 by lead compound Tenatoprazole (N16). The compound caused a dose-dependent shift in the T_m_ for Tsg101-UEV indicating binding to the key domain of Tsg101 as described in Materials and Methods.

We compared thermal denaturation profile for Tsg101-UEV in the presence and absence of tenatoprazole and found that it destabilizes the native protein structure, indicating that it binds Tsg101-UEV (Fig. 1). We also tested tenatoprazole against proteins unrelated to Tsg101, including DHPH, ENO1, MEK4, and did not observe a T_m_ shift (data not shown), indicating that the T_m_ shift of Tsg101-UEV was due to specific interaction of the prazole compound. This specific binding is consistent with a previous NMR structure in which tenatoprazole forms a covalent disulfide bond to Cys73 in the UEV domain of the protein (5). This disulfide bond formation can be prevented by including the reducing agent DTT in the assay (Fig. S1). The addition of DTT abolished the Tsg101-UEV T_m_ shift caused by the prazoles. Therefore, the addition of DTT to the FTS assay is a facile means to ascertain if prazole analogs interact with Tsg101-UEV in a covalent manner.

### Tenatoprazole inhibits herpes virus release from Vero cells

Tenatoprazole and esomeprazole were shown to quantitatively inhibit the release of infectious HIV-1 from 293T cells in culture, and it was suggested that these effects may be mediated via changes in viral interaction with Tsg101, a key component of the cellular ESCRT complex (5, 33). Given multiple reports suggesting that herpes viruses also use cellular ESCRT proteins in their replication process (20–23) we tested if the Tsg101-binding prazole drugs, which blocked budding of HIV-1, would also block the release of herpes viruses from cells.

We infected Vero cells with HSV-1 and HSV-2 for two hours at a multiplicity of infection (MOI) of 0.1 to assay the antiviral activity of tenatoprazole. Following infection, cells were treated with different concentrations of tenatoprazole. After 24 or 48 h the media fractions were collected and released virus titers were determined by standard plaque assays (34). Tenatoprazole caused a 3-log drop of HSV-1 and 4-5 log drop of HSV-2 of released virus titer from Vero cells in a dose dependent manner (Table 1, column 2, 3, and 5) with calculated EC_50_’s ranging from 48-80 μM. Total virus titer was also determined to differentiate between virus released into the media and infectious particles present in cell lysate. Total infectious virus particles were also reduced by tenatoprazole (Table 1, column 4). The concentrations of tenatoprazole that blocked virus release were nontoxic to Vero cells as determined by a 96^®^ AQueous One Solution cell proliferation assay reagent (Table 1, column 6). Taken together, tenatoprazole inhibited levels of both released and infectious virus particles without affecting cell viability at effective concentrations.

**Table 1.** Effect of Tenatoprazole on HSV-1 and-HSV-2 release from Vero cells. Tenatoprazole was incubated with Vero cells infected with HSV-1 or HSV-2 at a range of concentrations. The virus released into the media fraction at stated times was determined as described in Materials and Methods. Total virus is the amount of virus released from cells plus virus inside of the cells. Viability of Vero cells incubated with increased concentration of tenatoprazole was determined using the 96^®^ AQueous One Solution cell proliferation assay reagent as described in Materials and Methods.

| Tenatoprazole<br>(μM) | Titer of HSV-1<br>Media, 24 h | Titer of HSV-2<br>Media, 24 h | Total Titer HSV-2<br>Media + Cell<br>Lysate, 24 h | Titer of HSV-2<br>Media, 48 h | Viability<br>OD 490nm, 24 h |
| --- | --- | --- | --- | --- | --- |
| 0 | 2.50E+05 | 2.80E+05 | 4.70E+07 | 8.50E+06 | 1.694 |
| 52 | 2.90E+05 | 6.50E+04 | 4.00E+06 | 1.30E+06 | 1.724 |
| 60 | N.D. | 1.00E+03 | 2.30E+04 | 5.60E+04 | 1.759 |
| 79 | 1.30E+05 | 2.50E+02 | 1.50E+03 | 4.80E+03 | 1.742 |
| 105 | 5.40E+04 | 0.00E+00 | 6.00E+02 | 1.80E+02 | 1.777 |
| 131 | 2.40E+03 | 0.00E+00 | 3.00E+02 | N.D. | 1.714 |
| 157 | 1.30E+02 | N.D. | N.D. | N.D. | N.D. |
| 200 | N.D. | N.D. | N.D. | N.D. | 0.872 |

### Cellular location of tenatoprazole inhibition

We next imaged herpes virus infected-Vero cells using transmission electron microscopy to determine the site of inhibition of release of virus and whether it was similar to observations of HIV-1 release from 293T cells. Vero cells grown on glass cover slips were infected with HSV-2 at MOI of 0.1 for 2 h and then treated for 24 h with 105 μM tenatoprazole or vehicle control. Using electron microscopy, we examined eighty cells with virus particles, and representative images are shown in Fig. 2. In the control, virus particles were in both the nucleus and cytoplasm near the cell surface (Fig. 2A). In the tenatoprazole-treated cells the cytosol of the intact cells was largely devoid of virus particles (Fig. 2B). Instead, we observed large pockets of granular material accumulated in the nucleus and immature virus particles lined on the inside of the nuclear membrane (inset, B). These results suggest that tenatoprazole blocks the passage of herpes virus particles through the nuclear membrane. This result is different from that observed with HIV-1. Because tenatoprazole binds Tsg101, it suggests that the ESCRT-I protein complex is involved in transport of HSV-2 through the nuclear membrane.

**Figure 2.**
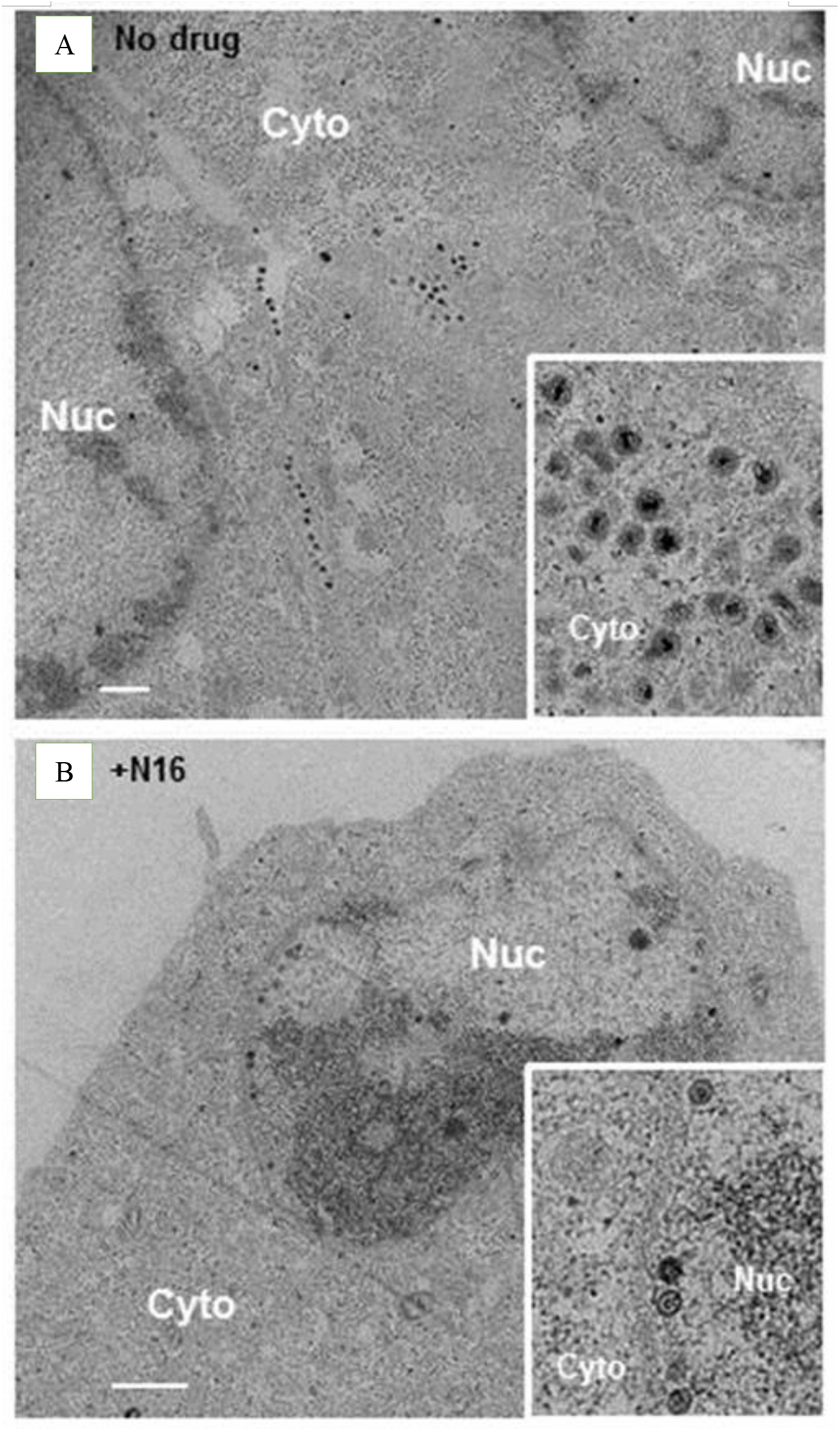
Inhibitory effect of tenatoprazole on HSV-2 production and location of virus particles inside of infected cells. Cells with virus were untreated (A) or treated with 105 μM tenatoprazole (N16) (B) for 24 h and examined by transmission electron microscopy. In each case, 80 cells where virus particles were observed were examined. Bar = 1 μm. *Inset*, higher magnification image.

### Identification of potent prazole-based inhibitors

Despite the lack of cell toxicity signal at effective tenatoprazole concentrations, the effective concentration is too high for use as a clinical therapy. Therefore, more potent analogs are required to explore antiviral therapeutic potential. We set out to identify and test other analogs which were more potent prazole analogs. We began by searching PubChem for analogs of tenatoprazole. We identified and obtained a dozen such compounds from commercial sources and prioritized these for testing based on structural similarities around the sites where tenatoprazole covalently linked to Cys73 of Tsg101. To this end, tenatoprazole, lansoprazole, rabeprazole, dexlansoprazole, pantoprazole, esomeprazole, 4-desmethoxy-omeprazole (an omeprazole analog, 5-methoxy-2[[(3,5-dimethyl-4-methoxy-pridin-2-yl-N-oxide)methyl]sulfinyl]-1H-benzimidazole; labelled O-Omeprazole), omeprazole, and ilaprazole were assessed in the FTS assay for their ability to change the T_m_ of Tsg101-UEV as described above (data not shown).

We determined the dose response plots of Tsg101 melting temperature shifts caused by these prazole compounds binding to Tsg101 (1-145) (Fig. 3). O-omeprazole is the only compound predicted not to form the covalent bond with Tsg101, since it has an oxygen linked to a ring nitrogen that is normally a hydrogen in the active prazoles (Table 2, right column). As expected, O-omeprazole did not cause a detectable thermal shift. The smallest thermal shift was observed with pantoprazole (gray) and the largest thermal shift was observed with ilaprazole (green). Ilaprazole’s ability to cause a thermal shift with Tsg101 was blocked by the addition of DTT (Fig. S1), consistent with the idea that the compound forms a disulfide linkage to Tsg101.

**Figure 3.**
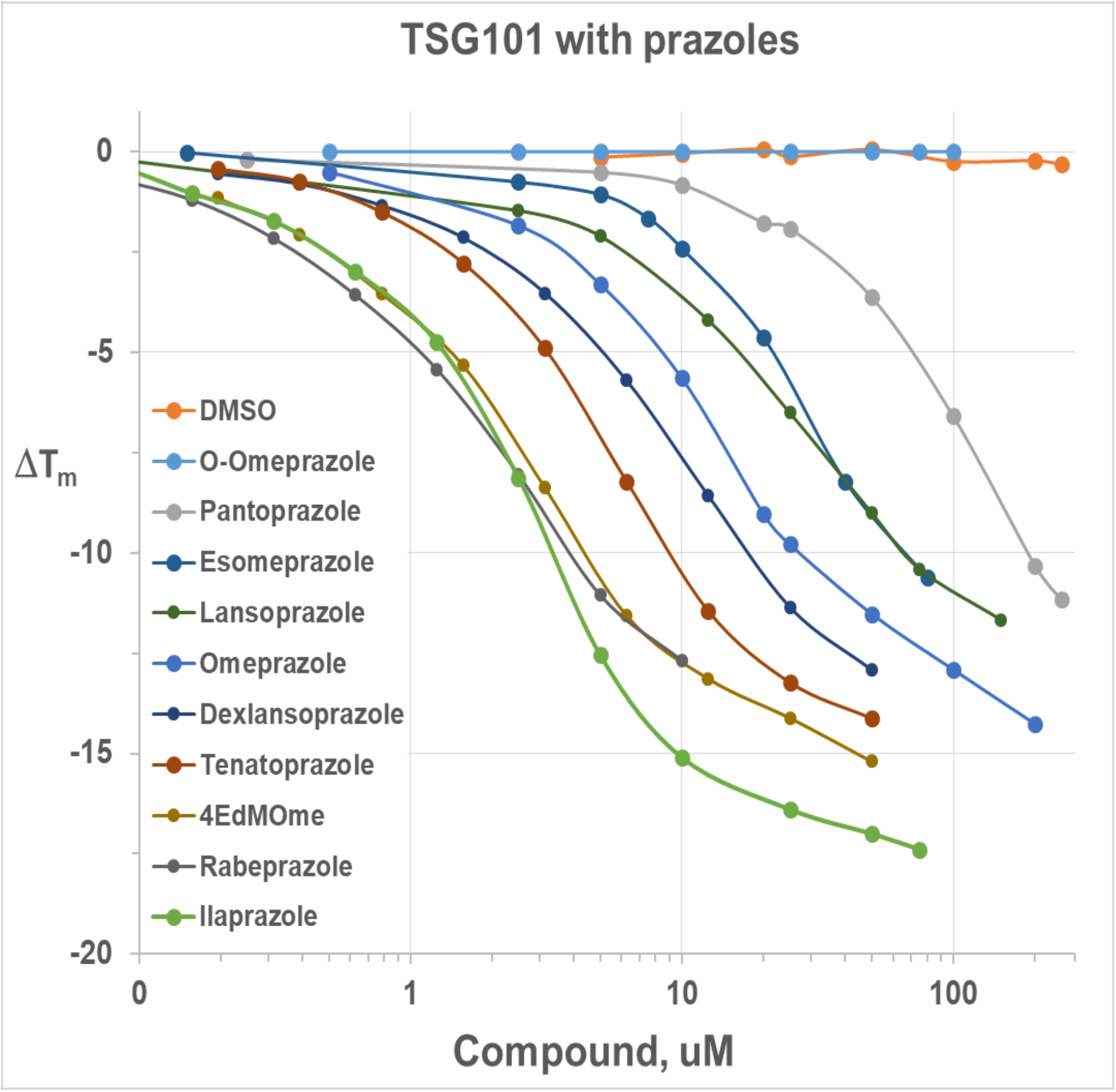
Dose-response plots of Tsg101 melting temperature shift caused by 10 prazole compounds. Different concentrations of prazole compounds were incubated with Tsg101 (aa 1-145) and subjected to Fluoresecent Thermal Shift analysis as described in Materials and Methods.

**Table 2.** Effect of prazole analogs to inhibit the release of HSV-2 from Vero cells. Different concentrations of the listed prazole drugs were incubated with HSV-2 infected Vero cells for 24 hours and then virus released into the media was quantified by plaque assays. Data presented includes the EC_50_ value (concentration at which virus release is inhibited by 50%). Methods are as described in the legend of Table 1.

| Prazole Compounds | EC<sub>50</sub> (μM)<br>Inhibition of HSV-2<br>Budding at 24h | Structure |
| --- | --- | --- |
| O-omeprazole | - |  |
| Pantoprazole | - |  |
| Esomeprazole | 140 |  |
| Lansoprazole | 84 |  |
| Omeprazole | 78 |  |
| Dexlansoprazole | 76 |  |
| Tenatoprazole | 84 |  |
| 4-Desmethoxy-omeprazole | 52 |  |
| Rabeprazole | - |  |
| Ilaprazole | 3-9 |  |

Next, we tested the anti-herpes virus activity of these prazole compounds (Table 2). To examine the effects of the compounds on the release of HSV-2 from Vero cells, we infected the cells with virus two hours prior to treatment with media containing different concentrations of compound. We incubated the cells for 24 or 48 hours and then collected the cell media fractions. Several of the analogs were inactive, including O-Omeprazole, pantoprazole, and rabeprazole. We identified a number of active compounds, in which there was a 10-fold spread of inhibition activity against HSV-2, ranging from an EC_50_ of 140 μM (for esomeprazole) to 3-9 μM (for ilaprazole). Thus, we identified prazole analogs that are more potent than tenatoprazole.

We provide the structures of prazole compounds tested in this analysis (Table 2, column 3). Of note, ilaprazole contains an additional ring structure compared to tenatoprazole that is predicted to lie in a solvent exposed area of the Tsg101 structure that may serve to strengthen the interaction with Tsg101. In examining the thermal shift capacity of the prazoles, we found that these roughly correlated to their HSV-2 antiviral activity. This correlation indicates that the FTS assay is useful in evaluating structure-activity-relationships (SAR) to inform the design of new compounds (Fig 3, Table 2).

### Antiviral activity of Ilaprazole on HSV-1, HSV-2, and HIV-1 in vitro

Based on these HSV-2 antiviral assay results, we selected ilaprazole for further antiviral profiling. First, we tested it against HSV-1 (Table 3, columns 2-4). Ilaprazole was slightly more effective against HSV-1 than against HSV-2 with EC_50_ calculations ranging from 0.6-5 μM (Table 3). Ilaprazole’s potency is a large improvement over tenatoprazole, which inhibited in the high μM range (Table 1 & 3). Additionally, ilaprazole was even more effective in inhibiting virus release at 72 h as at 24 h after a single application of the drug (72 h EC_50_<1μM; compare Table 3, columns 2 & 4). The inhibition caused by tenatoprazole against either virus began to fall off after 48 h (data not shown). We also tested for toxicity in the range of effective concentrations and did not observe cell toxicity using the 96^®^ AQueous One Solution cell proliferation assay reagent and WST-1 reagent over a 24 h period (Table 3, right columns). Thus, ilaprazole is more potent and has longer lasting effects than tenatoprazole.

**Table 3.** Effect of Ilaprazole on release of HSV-1 and HSV-2 from Vero cells. Different concentrations of ilaprazole were incubated for the times indicated with HSV-1 or HSV-2 infected cells similar to that described in the legend to Table 1. Virus titer released into the media and total virus was determined. Viability of Vero cells incubated with increased concentration of tenatoprazole was determined using the 96^®^ AQueous One Solution cell proliferation assay reagent as described in Materials and Methods.

| Ilaprazole (μM) | Titer of HSV-1 in media, 24h | Titer of HSV-1 in media, 48h | Titer of HSV-1 in media, 72h | Titer of HSV-2 in media, 24h | Total HSV-2 in media + cell lysate, 24h | Titer of HSV-2 in media, 48h | Ilaprazole (μM) | 96® Cell Viability OD<sub>490nm</sub>, 24h | WST-1 Cell Toxicity OD<sub>440-660nm</sub>, 24h |
| --- | --- | --- | --- | --- | --- | --- | --- | --- | --- |
| 0 | 3.00E+06 | 3.90E+07 | 1.00E+08 | 2.80E+05 | 1.20E+07 | 1.00E+06 | 0 | 1.694 | 0.993 |
| 4.5 | 2.00E+06 | 2.40E+06 | 9.00E+05 | 1.00E+05 | 3.60E+07 | 2.50E+05 | 4.5 | 1.764 | 1.058 |
| 9.0 | 7.50E+04 | 2.50E+05 | 2.20E+05 | 5.00E+04 | 4.50E+06 | 7.50E+04 | 9.0 | 1.711 | 1.055 |
| 13.5 | 3.20E+04 | 2.00E+02 | 4.50E+02 | 1.00E+04 | 4.30E+06 | 5.50E+04 | 13.5 | 1.690 | 0.950 |
| 18.0 | 6.00E+02 | 0.00E+00 | 1.00E+02 | 1.50E+03 | 2.00E+05 | 1.50E+03 | 18.0 | 1.737 | N.D. |
| 22.5 | 1.00E+02 | 0.00E+00 | N.D. | 1.00E+02 | 1.50E+04 | 3.00E+02 | 27 | N.D. | 1.055 |
| 54 | N.D. | N.D. | N.D. | N.D. | N.D. | N.D. | 54 | 1.658 | N.D. |
| 270 | N.D. | N.D. | N.D. | N.D. | N.D. | N.D. | 270 | 0.466 | 0.423 |

We next carried out a transmission electron microscopic examination of cells infected with HSV-1 at a MOI 0.1 in the presence and absence of 18 μM ilaprazole to determine if this drug causes the accumulation of virus particles in the nucleus of cells similar to tenatoprazole. Without drug, we observe particles in the cytoplasm and in the nucleus (Fig 4A & C), in the presence of drug little or no viral particles are found in the cytoplasm (Fig 4B & D). In both heavily infected cells (Fig 4A & B) and mildly infected cells (Fig 4C & D), treatment lead to particles being detected in the nucleus arrayed along the nuclear membrane (Fig 4C & D). This indicates that location of particles in the cell in the presence of drug is independent of the number of particles observed. These results are similar to the effect of tenatoprazole on HSV-2 infected cells (Fig. 2).

**Figure 4.**
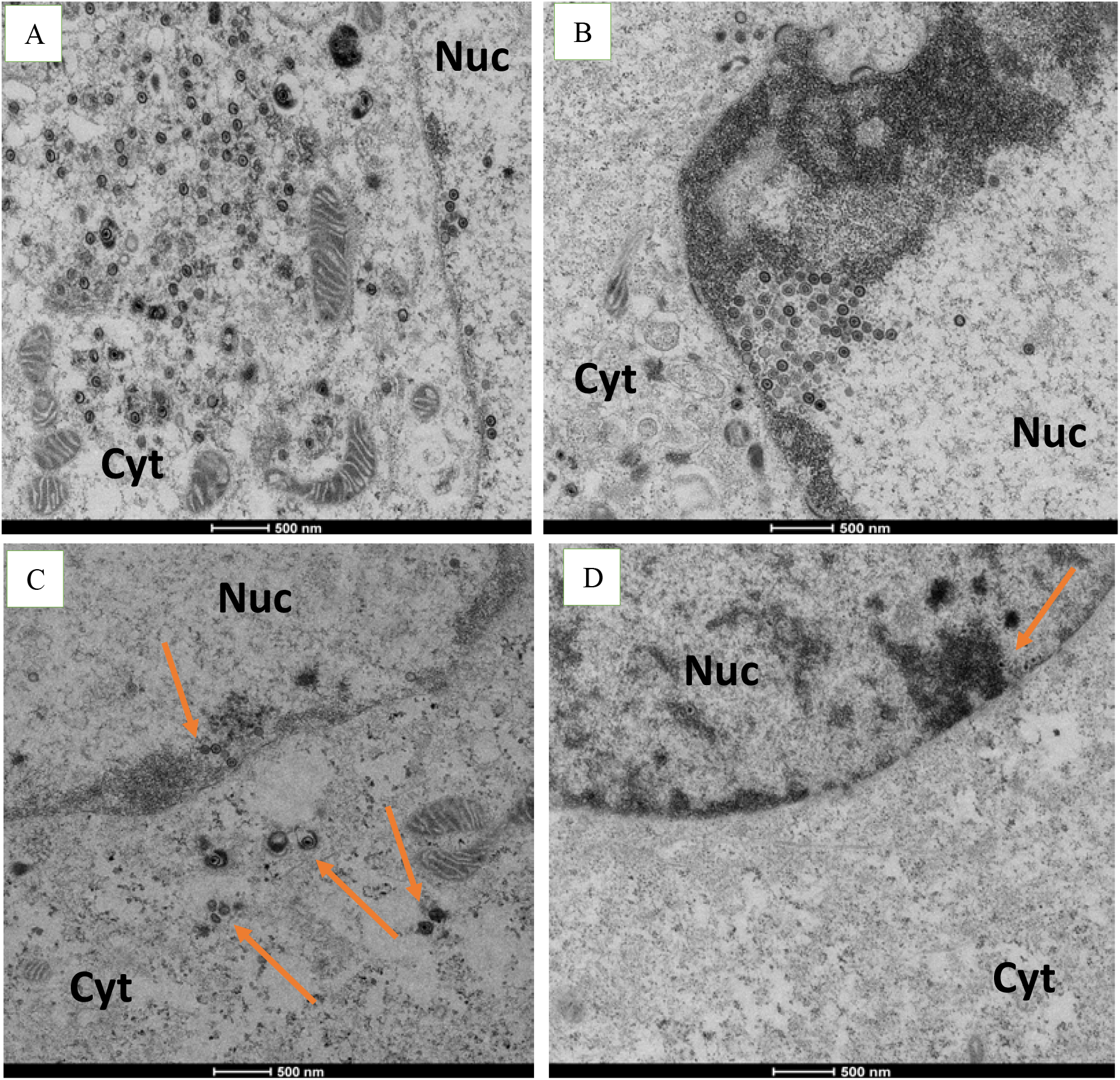
Inhibitory effect of Ilaprazole on HSV-1 production. Vero cells infected with HSV-2 at MOI of 0.1 and examined by transmission electron microscopy 24 h later. (A & C) untreated cells. (B & D) cells treated with ilaprazole (18 μM). Eighty cells where virus particles were observed were examined. In panels C and D, arrows point to virus particles. Nuc, nucleus. Cyt, cytoplasm.

Finally, to confirm the broad-spectrum potential of ilaprazole, we tested whether ilaprazole would inhibit the release of HIV-1 from 293T cells. To this end, cells were transfected with pR9-HIV-1_Ba-L_ plasmid and release of virus into the media fraction was detected by monitoring the capsid (CA) protein (p24) via enzyme linked immunosorbent assay analysis (ELISA). The drug was tested at concentrations between 0 and 40 μM and the effect of the drug on release of virus assessed (Table 4). Ilaprazole was effective at inhibiting the release of HIV-1 from cells with a calculated EC_50_ of 0.8 μM. We did not detect toxicity to the cells at the drug concentrations that inhibited the release of HIV-1 over the course of these experiments (data not shown).

**Table 4.** Effect of ilaprazole and novel analogs on release of HIV from 293T cells. Different concentrations of ilaprazole and novel compounds were incubated with HIV-1 infected cells as described in Materials and Methods. Virus titer released into the media was determined by monitoring p24 levels in the media 24 h post-infection.

| Ilaprazole<br>μM | p24 pg/ml<br>HIV | NWU-10<br>μM | p24 pg/ml<br>AVG<br>HIV | NWU-29<br>μM | p24 pg/ml<br>HIV |
| --- | --- | --- | --- | --- | --- |
| 0 | 7021 | 0 | 7021 | 0 | 7021 |
| 0.5 | 4063 | 0.5 | 1216 | 0.5 | 1197 |
| 1 | 3518 | 1 | 1080 | 1 | 1127 |
| 2 | 1652 | 3 | 663 | 3 | 787 |
| 5 | 274 | 5 | 448 | 5 | 518 |
| 10 | 178 | 10 | 411 | 10 | 303 |

### Identification of a novel prazole-based viral budding inhibitors

In an effort to design and identify more potent viral budding inhibitors, we synthesized 53 novel prazole-based analog compounds and assessed these for binding to Tsg101-UEV using FTS. Of the compounds screened, eight compounds demonstrated a T_m_ shift greater than or equivalent to that observed with ilaprazole (selected examples, Fig 5) indicating that these compounds bind to Tsg101. Based on the results described above, we are currently assessing the antiviral activity of these compounds in multiple antiviral assays. Initial testing of four analog compounds against HIV-1 reveal further increases in potency above that seen with ilaprazole (Table 4) with EC_50_ calculations ranging from 14-16 nM. This supports the pursuit of a medicinal chemistry campaign to apply the SAR learned from the prazole drugs to identify and develop potent compounds with broad-spectrum antiviral potential.

**Figure 5.**
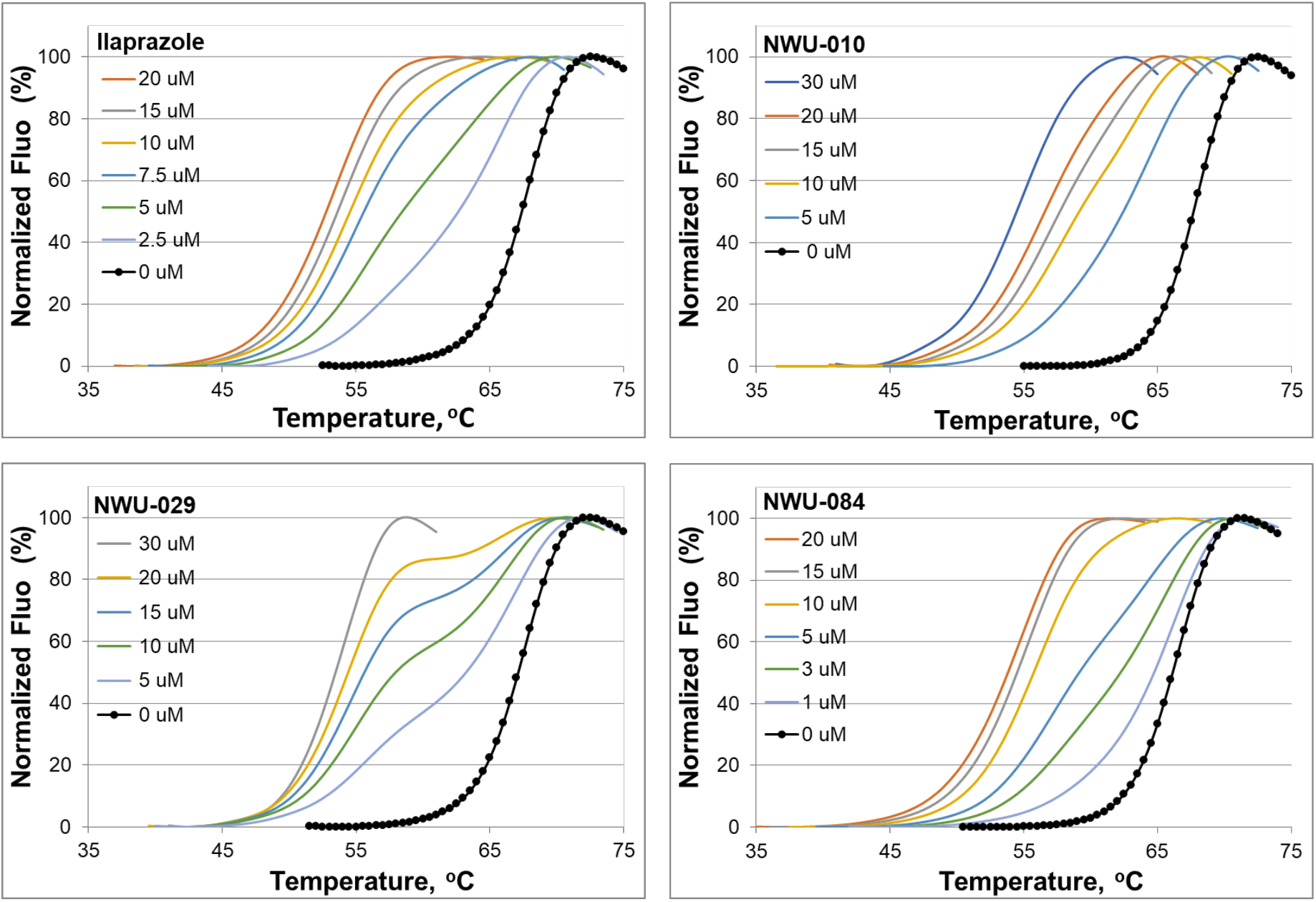
Thermal shift data of Tsg101 by lead compounds ilaprazole and select novel compounds. These compounds shifted the T_m_ for Tsg101-UEV indicating binding to the key domain of Tsg101.

Taken together, our results indicate that prazole-based drugs block release of HIV-1 and herpes viruses (HSV-1 and HSV-2), two families of viruses with different replication mechanisms that share the common pathway of Tsg101-mediated release of virus particles. Interestingly, we were able to identify more potent prazole analogs, in particular ilaprazole as well as novel compounds that are now in development. We found that ilaprazole and our analogs demonstrated antiviral effects significantly more potent, and potentially longer lasting, than other prazoles we tested.

## Discussion

We are developing a novel strategy to treat viral infections affecting humans by disrupting a common mechanism used by many enveloped viruses to bud from cells. Viral budding Inhibitors (VBI) have the potential to be broad-spectrum antiviral therapeutics, potentially being effective against herpes (20–23, 35), retro/lenti- (5), arena- (LFV, LCMV) (36, 37), flavi- (HCV) (38, 39), filo- (Ebola, MarV) (40–47), hepadnavirus (HBV) (48), some paramyxoviruses (SV5, MuV) (49–51) and rhabdoviruses (VSV, RV) (9, 52, 53). VBIs would require testing for antiviral activity towards these different viruses before clinical use but nonetheless present a strong starting point for identifying therapeutics.

In this work we demonstrate antiviral activity of prazole compounds, with no detectable cell toxicity at effective concentrations, against two viruses that use different mechanisms for viral replication. Of particular note is that the viral genomes are very different, with HIV being RNA-based and HSV being DNA-based. That one compound works against viruses with such stark difference in viral life cycle types supports that these compounds have potential as a broad-spectrum antiviral agent for current and emerging viruses. This aspect gives this approach advantage over other potential broad-spectrum antivirals, such as remdesivir, which is targeted to RNA viruses, limiting its potential as a broadspectrum antiviral (54).

Tsg101 binding to the proline-rich PTAPP viral L-domains in Gag (3, 6, 7, 11, 14, 15) is required for virus particles to be released from cell membranes of infected cells. Tsg101 is a member of the ESCRT-I complex of proteins involved in cell endosomal sorting. The ESCRT-I complex recruits proteins from the ESCRT-III complex with AIP1 (19), which provides the mechanical means for scission of virus particles from cell membranes. Thus, blocking the PTAPP L-domain sequence from interacting with the host proteins causes the virus budding defect and several lines of independent evidence support this idea. First, drugs targeted to this specific interaction in HIV cause budding defects in cells without detectable off-target effects (5). Second, a research group identified noncoding SNPs in the 5’ region of Tsg101, which affect the degree of Tsg101-mediated release of viral particles (27). Third, viral infections activate a host innate immunity mechanism, through Interferon Stimulated Gene 15 (ISG15), that specifically disrupts virus budding complexes (1). In response to this immune system defense, many viruses encode enzymes that prevent or reverse ISG15 conjugation to cellular proteins to avoid the budding blockade (55–60).

While the prazole analogs block the release of HIV-1, HSV-1, and HSV-2, the inhibition is manifested in different areas of the cell. The drugs block the release of HIV-1 at the outer cell membrane by preventing pinching of virus particles from the membrane (5). In contrast, herpes viruses appear to be first blocked at the passage of the virus through the nuclear membrane. Because the prazole drugs form a covalent bond to Tsg101, it strongly suggests that the ESCRT proteins are important for the herpes virus particles to be released from the nucleus of the cell where they are formed. This is consistent with the recent report by Arii *et al*., (61) that the ESCRT-III protein complex mediates herpes virus movement across the nuclear membrane and regulates its integrity. The finding that the prazole drugs cause a significant drop in total infectious herpes viruses can be explained by the trapping of immature particles in the nucleus, which prevents them from migrating into the cytoplasm to exchange enveloped membranes, which makes them infectious. The accumulation of the dense material in the nucleus observed in the electron micrographs suggests that the drug may interfere with normal particle assembly in addition to blocking the release of the particles from the nuclear membrane.

The use of prazoles as an antiviral represents an exciting potential case of repurposing existing drugs to act as antiviral agents. Currently, omeprazole is marketed as a prodrug for treatment of acid reflux disease. The prodrugs are acid-activated into derivatives that form disulfide linkages with proton pumps (28, 62, 63). The prodrug, but not the charged sulfonamide derivative, can cross the plasma membrane barrier. The antiviral activity of tenatoprazole has been suggested to be the result of forming a covalent disulfide bond with Tsg101 (5). While the binding site for tenatoprazole is near the ubiquitin (Ub) binding pocket and not the L-domain binding site, biochemical and confocal imaging data independently demonstrated that tenatoprazole disrupts the binding of Tsg101 to the PTAPP sequence (5). While the precise biochemical mechanism remains to be clarified, our FTS results support that it may be related to allosteric changes in Tsg101 after the drug forms its covalent linkage with Cys73. One of our most potent lead compounds is another prazole drug, ilaprazole, which is also marketed for treatment of acid reflux disease in China, India, and South Korea (Yi Li An, Adiza, Noltec, respectively) indicating that it has reasonable bioavailability and a known clinical safety profile. Previous reports did not detect off-target effects of the prazole drugs affecting Tsg101 metabolism inside of cells (5). The prazoles we tested here also appear to be nontoxic to Vero, Hela, and 293T cells at the concentrations used to inhibit budding of herpes viruses and HIV-1.

A recent report highlighted the potential of prazole compounds to have a therapeutic effect on SARS-CoV-2 when combined with remdesivir (30). However, the authors did not definitively identify the mechanism of action of the prazoles and also concluded that the potency of the prazole compound used, omeprazole, is too low to reach therapeutic levels in vivo. A potential mechanism posed by the authors is that the prazoles lead to an increase in lysosomal pH, which is the potential mechanism for lysosomotropic drugs such as chloroquine (64). In contrast to omeprazole, we hypothesize that ilaprazole and our more potent novel prazole compounds may allow for therapeutic levels to be reached in vivo. In the case of ilaprazole, which is marketed in several Asian countries as discussed above, our strong in vitro results lay the foundation for a potential fast-track to broad-spectrum antiviral clinical testing, alone or in combination with other drugs, in these countries. We are currently working to determine if ilaprazole or our novel compounds have activity against SARS-CoV-2 with or in combination with remdesivir. This would further the potential broad-spectrum antiviral capacity of the prazole compounds described in this report.

## Materials and Methods

### Viruses, plasmids, cell lines

Herpes simplex virus-1 (Kos strain), Herpes simplex virus-2(A/B-G), HIV plasmid pNL4 (from Carol Carter), pET-28b vector (Novagen-EMD Millipore), ROSETTA 2 (DE3) pLysS *E. coli* competent cells (EMD Millipore), Vero cells and 293T cell lines.

### Chemicals

Prazole Compounds: Rabeprazole, Lansoprazole, Omeprazole, Ilaprazole, Dexlansoprazole, Tenatoprazole, and Pantoprazole were from SelleckChem. 2-[(4-ethoxy-3-methylpridin-2-yl)methanesulfinyl]-1H-1,3benzodiazole, 2-[(3,5-dimethylpyridin-2-yl)methanesulfinyl]-5-methoxy-1H-1,3-benzodiazole,4-methoxy-2-[[(5-methoxy-1H-1,3-benzodiazol-2-yl)sulfinyl] methyl]-3,5-dimethyl-1λ-pyridin-1-one were from MolPort. Esomeprazole was from Toronto Research Chemicals.

### Purification of Tsg101 (1-145)

N-terminally His6-tagged Tsg101 UEV domain (amino acids 1-145), called Tsg101-UEV, was encoded in a pET-28b vector (Novagen – EMD Millipore), which also included a thrombin protease cleavage site (His6-Thrombin Site-Tsg101, 1-145). Tsg101-UEV was grown in LB broth with Kanamycin (30 μg/ml) in ROSETTA 2 (DE3) pLysS *E. coli* competent cells (EMD Millipore) and induced with 1 mM IPTG at room temperature for 3 h. Bacteria were collected by centrifugation at 4,000 rpm for 10 min at 4°C. Bacteria were suspended in 50 ml binding buffer (20 mM Tris-HCl, pH 7.9, 0.5 M NaCl, 5 mM Imidazole) with 1 mM PMSF, 0.1% NP40, and a Protease Inhibitor Cocktail Tablet (Roche) and sonicated for 3.5 min on ice. The sonicate was spun at 9,000 rpm for 1 h at 4°C in a Sorvall centrifuge. The supernatant was collected and passed through a 1.5 ml Ni-NTA Agarose column. The column was washed with 20 mM Tris-HCl, pH 7.9, 0.5 M NaCl, 30 mM Imidazole wash buffer. The column was then equilibrated with TEV cleavage buffer followed by 50 units of thrombin in the same buffer (Novagene). The column flow was stopped and incubated at room temperature overnight. The cleaved protein was eluted with wash buffer, and the protein dialyzed in D-tube Dialyzer Maxi, MWCO 12-14 kDa (Novagene) overnight against 0.15 M NaCl, 0.1 M HEPES, pH 7.5 buffer. The protein was concentrated in a MicroSep Advanced Centrifugal Device, 12-14 kDa exclusion (Pall) for 1 h at 1,300 rpm). Protein concentration was determined with a Nano Drop Spectrophotometer at 280 nM. When the His tag was not removed, the protein was eluted from the Ni-NTA column with 20 mM Tris-HCl, pH 7.9, 0.5 M NaCl, 1 M Imidazole. The protein was evaluated by SDS-PAGE gel for purity.

### Fluorescence thermal shift (FTS) screening to identify small molecule binding to Tsg101-UEV

FTS using thermal shift elicited by the small molecule binding effect to protein stability. FTS monitors protein thermal denaturation using environment-sensitive dye Sypro^®^ Orange which fluoresces when bound to hydrophobic surfaces, taking advantage of the changes in hydrophobic surface exposure in protein denaturation. Discovery of small molecule binding to target protein utilizes the observation that ligand binding affects protein thermal stability, and therefore can be detected through a shift in the protein’s thermal denaturation (melting) temperature (T_m_). We have employed FTS to reveal changes in thermodynamic properties of Tsg101 elicited by its interaction with a small molecule. The recombinant Tsg101 fragment (amino acids 1-145), prepared as described above in *Materials and Methods* (but without label) has a thermal unfolding profile suitable for using FTS as a primary screen assay in HTS. A fluorescence dye Sypro^®^ Orange (Invitrogen) was used for assay detection. The dye is excited at 473 nm and has a fluorescence emission at 610 nm. The dye binds to hydrophobic regions of a protein that are normally buried in a native protein structure. When a protein is unfolded, the dye interacts with exposed hydrophobic surfaces and the fluorescence intensity increases significantly over that observed in aqueous solution. The Tsg101 fragment was premixed at a concentration of 2 μM with a 5X concentration of Sypro^®^ Orange in HEPES buffer (100 mM HEPES, 150 mM NaCl, pH 7.5). Then 10 μl of the protein-dye mix was added to an assay plate and 10 to 50 nanoliters of compound, equal to 10 to 50 μM, were added with an acoustic transfer robot Echo550 (Labcyte, CA). The plate was shaken to ensure proper mixing, then sealed with an optical seal and centrifuged. The thermal scan was performed from 20 to 90°C with a temperature ramp rate of 0.5°C/min. Fluorescence was detected on a real-time PCR machine CFX384 (Bio-Rad Laboratories). Comparison of the thermal denaturation profile for Tsg101-UEV in the presence and absence of tenatoprazole and other prazoles revealed destabilization of the native protein structure, indicating that the compound interacted with Tsg101-UEV.

### Herpes virus infection of Vero cells

Vero cells (0.8 × 10^6^ cells/well of a 6-well plate) were infected with HSV-1 or HSV-2 at a MOI of 0.1 in DMEM with 1% serum for two hours in the C0_2_ incubator at 37°C. The cell supernatants were aspirated and replaced with 1 ml (24 h) or 2 ml (48 h and 72 h) of DMEM with 1% serum with DMSO or different concentrations of drug dissolved in DMSO. After 24 or 48 h incubation, the cell supernatant was collected and frozen at −80°C. Virus titer in the cell media fraction was determined by standard plaque assays where cell supernatants were serially diluted, added to Vero cells and incubated for 48 h after which cells were fixed and stained to count the plaques (22). For determination of total virus (extracellular + cytoplasmic), virus infected cells were incubated for 24 h with and without drug presence, then the plate of cells were subjected to 3 cycles of freeze/thawing (−80°C/37°C) 30 min each prior to collecting the supernatant after centrifugation for measurement of total virus titer. Virus titer was measured by standard plaque assay as above. In separate experiments, uninfected Vero cells were carried for 3 weeks in culture in the presence or absence of drugs (replaced every third day) and found to exhibit the same growth rate detected with a light microscope.

### HIV-1 infection of 293T cells

293T cells (American Type Culture Collection) were grown in a 24-well Clear Flat Bottom TC-treated Multiwell Cell Culture Plate using Dulbecco’s modified Eagle’s medium (Cellgro) containing fetal bovine serum (10%), 100 U/ml penicillin, 100 μg/ml streptomycin, and 292 *μ*g/ml l-glutamine (Cellgro). Cells were grown to 60-70% confluency at 37 °C and 5% CO_2_ prior to addition of drug treatment. Culture media was aspirated and replaced with media containing drug compound 7 hours prior to transfection of the plasmid encoding the HIV-1 genome. Transfection was done using reagent Polyethyleneimine (PEI, Polysciences). For production of virus particles, cells were transfected with pR9-HIV-1_Ba-L_ plasmid. After 24 h and 48 h, tissue culture media was collected and passed through a 0.45 micron filter. Virus released from cells was quantified by media-associated p24 determined by ELISA (PerkinElmer) and equivalent amounts of p24.

### Drug potency and cell toxicity

EC_50_ calculations were determined by using AAT Bioquest’s EC_50_ calculator. Cell toxicity at different concentrations of drugs as indicated was determine using the Cell Proliferation Reagent WST-1 (Roche Diagnostics) or cellular 96^®^ Aqueous One Reagent viability reagent according to manufacturer’s instructions. For 293T cells, the concentration of DMSO was 0.2% or less and assays were carried out with DMEM with 10% serum.

### Transmission electron Microscopy

Vero cells on glass cover slips were infected with HSV-2 at a MOI of 0.1 for two hours. Then 105 μM of tenatoprazole or 18 μM Ilaprazole was added and cells incubated for 24 hours. Tissue samples were fixed in 0.1 M sodium cacodylate buffer pH 7.3 containing 2% paraformaldehyde and 2.5% glutaraldehyde and post-fixed with 2% osmium tetroxide in unbuffered aqueous solution. The samples were rinsed with distilled water, en bloc stained with 3% uranyl acetate, rinsed with distilled water, dehydrated in ascending grades of ethanol, transitioned with propylene oxide, embedded in the resin mixture of Embed 812 kit and cured in a 60°C oven. Samples were sectioned on a Leica Ultracut UC6 ultramicrotome. 1 μm thick sections were collected and stained with Toluidine Blue O and 70 nm sections were collected on 200 mesh copper grids; thin sections were stained with uranyl acetate and Reynolds lead citrate. Transmission electron microscopy (TEM) was performed on an FEI Tecnai Spirit G2.

## Acknowledgments

Transmission electron microscopy was performed by Mr. Lennell Reynolds at the Northwestern University Center for Advanced Microscopy, generously supported by NCI CCSG P30 CA060553 awarded to the Robert H Lurie Comprehensive Cancer Center. We gratefully acknowledge the use of Dr. Richard Longnecker’s lab for the HSV studies and outstanding technical support from Nanette Sosmaiski. Activity of compounds against HIV-1 was performed in the Pathology Core of the Center for Aids Research supported by the NIH NIAID grant 1P30AI117943. We thank Michael McRaven and Edward Allen for their technical expertise. This work was funded in part by the Chicago Biomedical Consortium with support from the Searle Funds at The Chicago Community Trust. and by funds from the Northwestern Memorial Hospital Dixon Innovation Grant and a grant from the Campbell Foundation (JL). Patent applications based on this work have been filed.

## Supplemental Figure

**Fig. S1.**
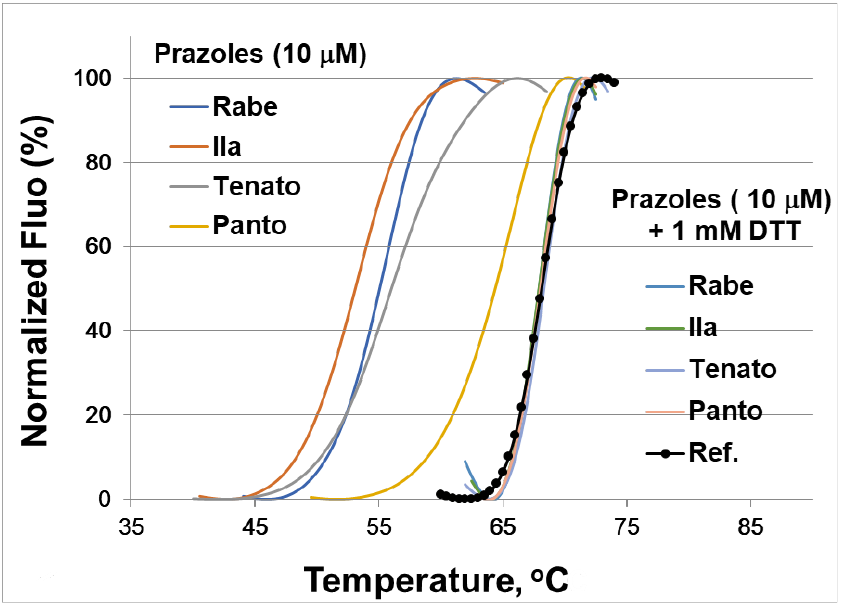
Effect of adding DTT to the thermal shift of Tsg101-UEV in the FTS assay. The addition of DTT abolishes the Tm shift in the FTS assay. This is consistent with all of these prazole compounds forming a disulfide bond to Tsg101.

## Notes

### Competing Interest Statement

The authors have declared no competing interest.

